# Global analysis of cell behavior and protein localization dynamics reveals region-specific functions for Shroom3 and N-cadherin during neural tube closure

**DOI:** 10.1101/2021.01.19.427312

**Authors:** Austin T. Baldwin, Juliana Kim, John B. Wallingford

## Abstract

Failures of neural tube closure are common and serious birth defects, yet we have a poor understanding of the interaction of genetics and cell biology during neural tube closure. Additionally, mutations that cause neural tube defects (NTDs) tend to affect anterior or posterior regions of the neural tube but rarely both, indicating a regional specificity to NTD genetics. To better understand the regional specificity of cell behaviors during neural tube closure, we analyzed the dynamic localization of actin and N-cadherin via high-resolution tissue-level time-lapse microscopy during *Xenopus* neural tube closure. To investigate the regionality of gene function, we generated mosaic mutations in *shroom3*, a key regulator or neural tube closure This approach elucidates new differences between cell behaviors during cranial/anterior and spinal/posterior neural tube closure, provides mechanistic insight into the function of *shroom3* and demonstrates the ability of tissue-level imaging and analysis to generate cell-biological mechanistic insights into neural tube closure.

## Introduction

Congenital birth defects are the number one biological cause of death for children in the US, and neural tube defects (NTDs) represent the second most common class of human birth defect (Murphy et al., 2018; Wallingford et al., 2013). NTDs represent a highly heterogenous group of congenital defects in which failure of the neural folds to elevate or fuse results in a failure of the skull or spine to enclose the central nervous system (Wallingford et al., 2013). While genetic analyses in both humans and animal models have revealed dozens of genes necessary for normal neural tube closure, several key questions remain.

One central unanswered question relates to the regional heterogeneity of both normal neural tube closure and pathological NTDs. For example, the collective cell movements of convergent extension dramatically elongate the hindbrain and spinal cord of vertebrates, but not the midbrain and forebrain (Nikolopoulou et al., 2017; Wallingford et al., 2013). Accordingly, disruption of genetic regulators of convergent extension such as the Planar Cell Polarity (PCP) genes results in failure of neural tube closure in posterior regions of the neural ectoderm, but not anterior (Kibar et al., 2001; Wang et al., 2006). Conversely, the Shroom3 gene is implicated in apical constriction, a distinct cell behavior that drives epithelial sheet bending, and disruption of *shroom3* elicits highly penetrant defects in anterior neural tube closure, but only weakly penetrant defects in the posterior (Haigo et al., 2003; Hildebrand and Soriano, 1999). This regional deployment of apical constriction in the anterior and convergent extension in the posterior during neural tube closure is poorly understood.

In addition, the underlying mechanisms of individual cell behaviors necessary for neural tube closure remain incompletely defined. While apical constriction is driven by actomyosin contraction, the precise site of actomyosin action during this process is unclear and constitutes a long-term problem in the field (Martin and Goldstein, 2014). For example, analysis of apical constriction during gastrulation in both *Drosophila* and *C. elegans* has shown integration of discrete junctional and medio-apical (“medial”) populations of actomyosin (Coravos and Martin, 2016; Martin et al., 2009; Roh-Johnson et al., 2012). Recent studies in frog and chick embryos have also described similar pulsed medial actomyosin-based contractions occurring during neural tube closure (Brown and García-García, 2018; Christodoulou and Skourides, 2015; Suzuki et al., 2017), but how those contractions are controlled and how they contribute to cell shape change during neural tube closure are not known.

For example, Shroom3 is among the more well-defined regulators of apical constriction, being both necessary and sufficient to drive this cell shape change in a variety of cell types, including the closing neural tube (Haigo et al., 2003; Hildebrand, 2005; Plageman et al., 2010; Plageman et al., 2011b). Shroom3 is known to act via Rho Kinase to drive apical actin assembly and myosin contraction (Das et al., 2014; Hildebrand, 2005; Nishimura and Takeichi, 2008; Plageman et al., 2011a). However, the relationships between Shroom3, and the medial and junctional populations of actin have not been explored.

An additional outstanding question relates to the interplay of actomyosin contractility and cell adhesion during apical constriction. The classical cadherin Cdh2 (N-cadherin) is essential for apical constriction during neural tube closure in *Xenopus* (Morita et al., 2010; Nandadasa et al., 2009), and *shroom3* displays robust genetic interactions with *n-cadherin* in multiple developmental processes, including neural tube closure (Plageman et al., 2011b). Moreover, a dominant-negative N-cadherin can disrupt the ability of ectopically-expressed Shroom3 to induce apical constriction in MDCK cells (Lang et al., 2014). Nonetheless, it is unclear if or how Shroom3 controls the interplay of N-cadherin and actomyosin during apical constriction. This is an important gap in our knowledge, because despite the tacit assumption that cadherins interact with each other and control actomyosin at cell-cell junctions, N-cadherin displays multiple cell-autonomous activities (Rebman et al., 2016; Sabatini et al., 2011). Intriguingly, several papers now demonstrate that extra-junctional cadherins at free cell membranes can engage and regulate the actomyosin cortex (Ichikawa et al., 2020; Padmanabhan et al., 2017; Sako et al., 1998; Wu et al., 2015).

Finally, though Shroom3 has been extensively studied in the context of apical constriction, recent studies also implicate Shroom family proteins in the control of convergent extension (McGreevy et al., 2015; Nishimura and Takeichi, 2008; Simões Sde et al., 2014). Several studies indicate a genetic and cell biological interplay of Shroom3 and the PCP proteins (Durbin et al., 2020; McGreevy et al., 2015), and one study directly links PCP, apical constriction and convergent extension (Nishimura et al., 2012). Conversely, some studies also suggest a role for PCP proteins in apical constriction (Ossipova et al., 2015). Given the better-understood role of PCP in regulating convergent extension in the posterior neural ectoderm, how Shroom3 controls convergent extension and PCP in the posterior remains especially unclear.

Together, these studies highlight the complexity of neural tube closure, which is compounded by the sheer scale of the tissue involved. The neural ectoderm is comprised of hundreds to thousands of cells (depending on organism) and stretches from the anterior to posterior poles of the developing embryo. However, the vast majority of dynamic studies of cell behavior in the neural tube closure, including our own, have focused on small numbers of cells due to constraints of both imaging and image analysis.

Here, we used image-tiling time-lapse confocal microscopy to obtain over 500,000 individual measurements of cell behaviors associated with neural tube closure in *Xenopus tropicalis*. Using these data, we demonstrate that the cell biological basis of apical constriction differs substantially between the anterior and the posterior neural plate. Second, we demonstrate that the crux of Shroom3 function lies not in actin assembly *per se*, but rather in the coupling of actin contraction to effective cell shape change. Third, we demonstrate that the control of N-cadherin localization is also a key feature of Shroom3 function during neural tube closure. Finally, we demonstrate that the incompletely penetrant posterior phenotype of *shroom3* mutants also results from dysregulation of both actin and N-cadherin localization. Overall, these findings *a)* elucidate differences between cell behaviors during cranial/anterior and spinal/posterior neural tube closure, *b)* provide mechanistic insight into the function of *shroom3*, an essential neural tube closure gene, and *c)* demonstrate the ability of tissue-level imaging and analysis to generate cell-biological mechanistic insights into neural tube closure.

## Results and Discussion

### High-content imaging of cell behavior and protein localization during vertebrate neural tube closure

*Xenopus tropicalis* affords several advantages for imaging neural tube closure, as its cells are large and easily accessible; its culture conditions for imaging are no more complex than synthetic pond water held at room temperature; and its broad molecular manipulability allows examination of diverse fluorescent markers. We used confocal microscopy and image tiling to collect high-magnification datasets spanning broad regions of the folding neural ectoderm from embryos injected at blastula stages with mRNAs encoding fluorescent reporters (**Fig. 1A**). At the onset of neurulation, embryos were positioned to image either the anterior (roughly corresponding to the brain) or the more posterior (roughly corresponding to the spine) regions of the neural ectoderm. Cells captured in the resulting movies were then segmented using Tissue Analyzer and CSML (Aigouy et al., 2016; Ota et al., 2018), yielding a map of both the apical cell surfaces and all individual junctions (**Fig. 1B**). Using Tissue Analyzer (Aigouy et al., 2010; Aigouy et al., 2016) and Fiji (Schindelin et al., 2012), we quantified both cell behaviors and the localization of fluorescent protein reporters.

**Figure 1:**
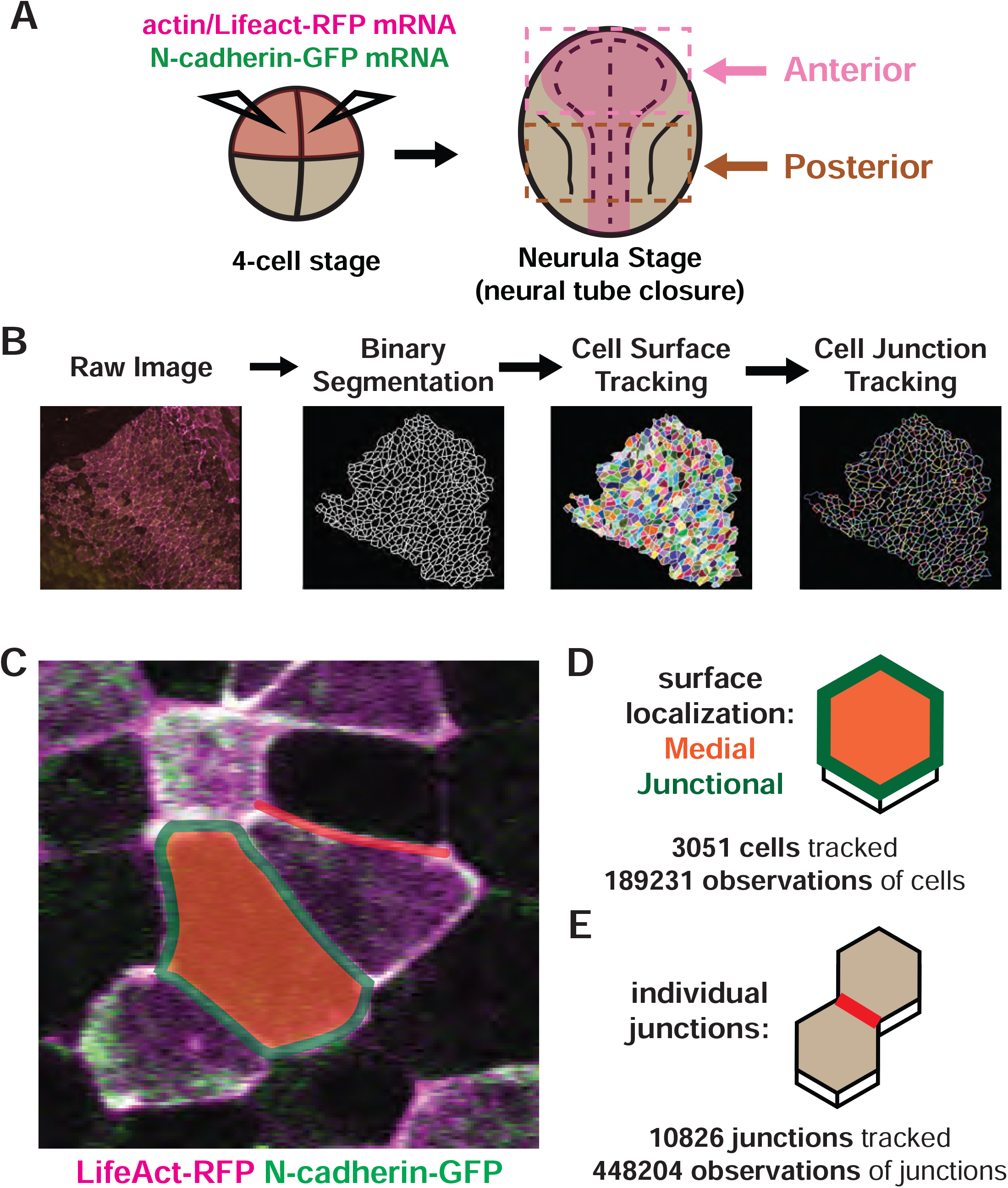
Tissue-level imaging and analysis of contractile protein dynamics during neural tube closure in *Xenopus*. **A**. Schematic of mRNA injections and subsequent imaged regions of the *Xenopus tropicalis* embryo. **B**. Cell segmentation and tracking workflow. Binary Segmentation, Cell Surface Tracking, and Cell Junction Tracking were all generated using Tissue Analyzer. **C**. Example *Xenopus* cells with analyzed subcellular domains labeled. Orange label = medial, green label = junctional, red label = individual junction. **D**. Schematic and N values of whole cell measurements. **E**. Schematic and N values of individual cell junction measurements.

Actomyosin contractility is the key driver of apical constriction, so we labeled actin using LifeAct-RFP (Riedl et al., 2008) to quantify actin localization (**Fig. 1A, magenta**). Because previous studies suggest that distinct actomyosin populations at the medial apical surface and the apical cell junctions play distinct roles in apical constriction (Coravos and Martin, 2016; Martin et al., 2009; Roh-Johnson et al., 2012), we quantified these populations separately (**Fig. 1C, D**). Because cell adhesion is also essential for epithelial morphogenesis and N-cadherin is essential for neural tube closure in *Xenopus* (Nandadasa et al., 2009), we co-labeled cells with N-cadherin-GFP (**Fig. 1A, green**).

In total, our dataset is comprised of ∼190,000 observations of apical cell surfaces from over 3000 cells and ∼450,000 observations from over 10,000 individual cell-cell junctions (**Fig. 1D, E**), collected at a rate of one frame/observation per minute over 1-2 hours across seven embryos spanning roughly stages 13 to 18 (Nieuwkoop and Faber, 1994).

As an initial test of the validity of our approach, we examined our dataset for well-known trends expected for neural epithelial cells during neural tube closure, namely an overall decrease in apical area and an overall increase in apical actin intensity. The heat maps in **Fig. 2A** and **B** reveal that cells in both anterior and posterior regions generally reduce their apical surface area and increase medial actin intensity, as expected. These overall trends are backed by examination of individual cells, as shown for specific representative cells in **Fig. 2C**. Overall, this analysis suggests that our pipeline is generally effective for quantifying cell shape and actin intensity in over time during neural tube closure.

**Figure 2:**
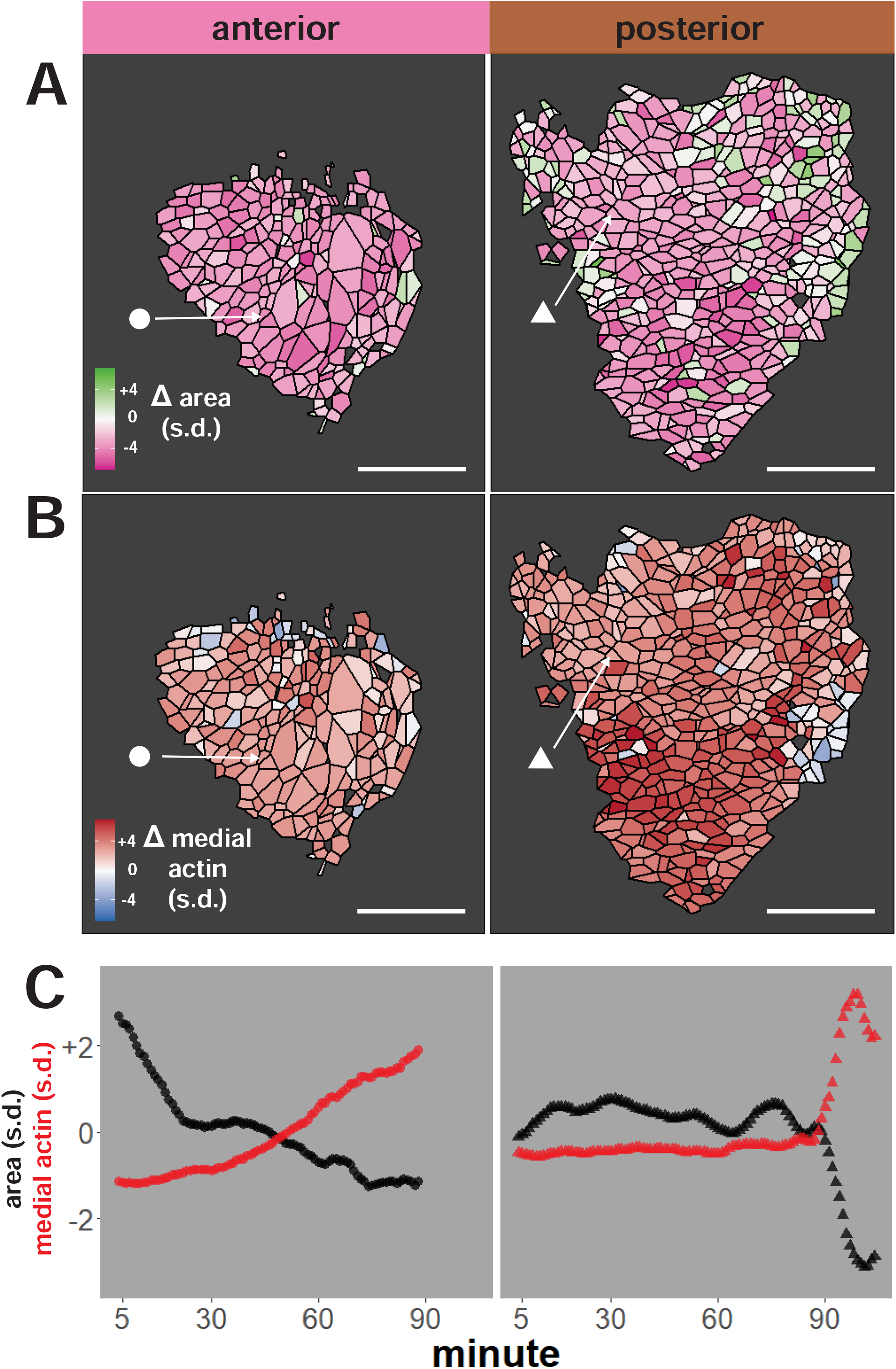
Tissue-level analysis of individual cell behaviors reveals dynamic heterogeneity. **A**. Overall change (Δ) in apical surface area (standardized) across anterior (left) and posterior (right) control embryos. **B**. Overall change in medial LifeAct/actin localization (standardized) across anterior (left) and posterior (right) control embryos. Circle and triangle in A & B denote a representative cell for each embryo. Scale bars = 100 microns. **C**. Standardized apical surface area (black) and medial actin (red) over time in representative cells from anterior (left/circle) and posterior (right/triangle). s.d. = standard deviation.

### Distinct patterns of behavior drive apical constriction in the anterior and posterior regions of the neural plate during neural tube closure

The incidence and form of NTDs differ between the brain and spinal cord in both humans and mouse genetic models (Nikolopoulou et al., 2017; Wallingford et al., 2013), yet how the dynamics of individual cell behaviors differ between the anterior and posterior regions remains poorly defined. We therefore quantified the temporal progression of apical constriction between the two regions, and we observed consistent and significant differences.

As noted above, bulk measurements show a decrease in apical area in both regions over time (**Fig. 3A**). However, we observed distinct region-specific distributions for these changes. For example, a significantly higher proportion of anterior cells displayed significant apical constriction as compared to cells of the posterior region (**Fig. 3A**). We also observed differences in the timing of constrictions, with anterior cells displaying a more gradual reduction in apical area across the period of observation, while cells in the posterior display late, rapid reductions in apical area (**Fig. 3A’**).

**Figure 3:**
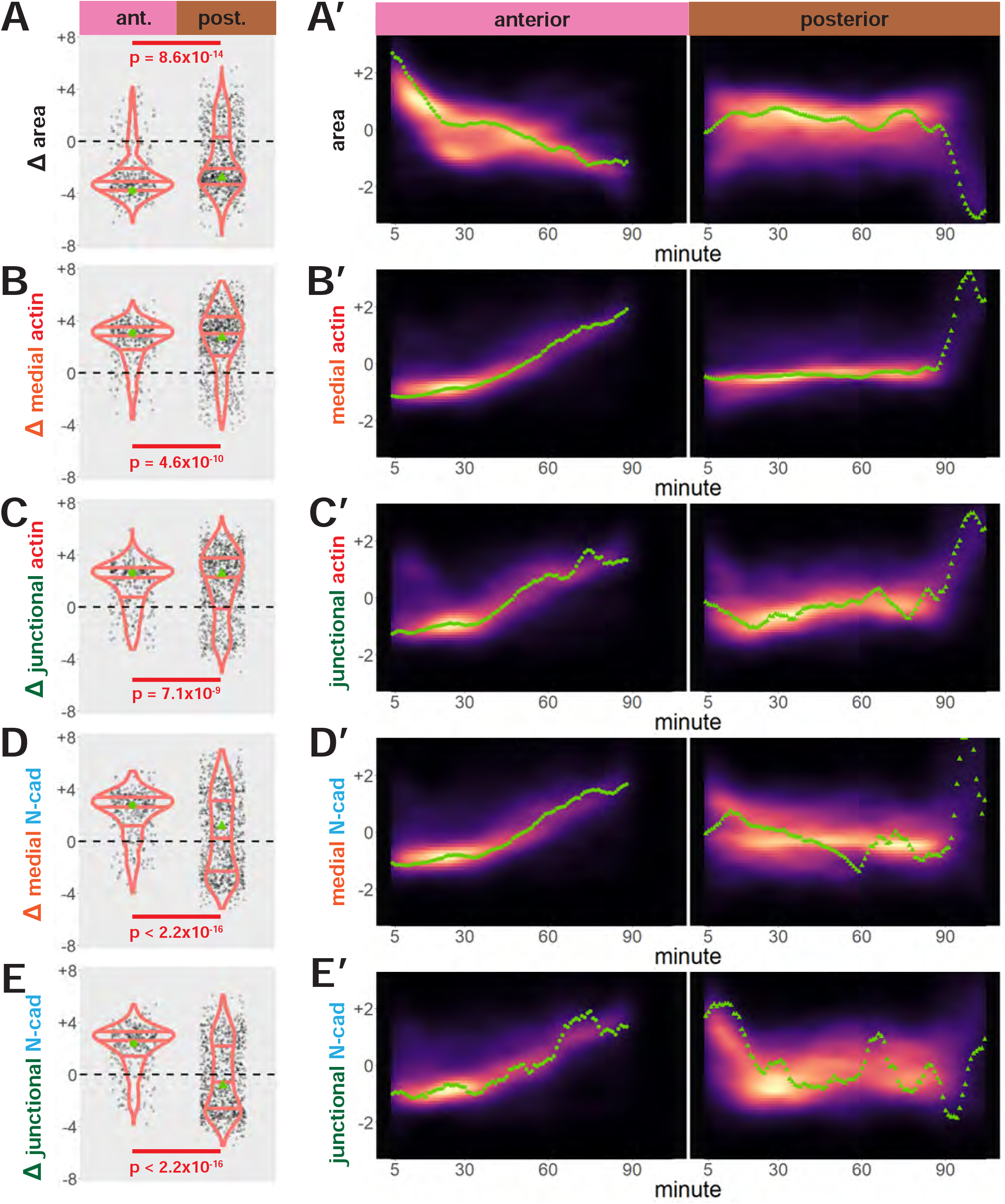
Cells in the anterior and posterior neural ectoderm both apically constrict but differ in their contractile protein dynamics. Tissue-level cell size and protein localization dynamics from control embryos in Figure 2. **X**. Distribution of overall change (Δ) in displayed parameter (standardized) among cells from control embryos. Horizontal lines on density plots/violins indicate quartiles of distribution. Black circles are individual cells. Statistical comparisons performed by Kolmogorov-Smirnov (KS) test. **X’**. 2D density plots of standardized variable versus time for all observations/cells in each control embryo in Figure 2. Green points are measurements from the representative cells denoted in Figure 2. s.d. = standard deviation.

Medial and junctional actin intensity also generally increased over time in both regions, with their temporal progressions reflecting the changes in apical area described above (**Fig. 3B, B’, C, C’)**. Likewise, the distributions of changes in both medial and junctional actin were significantly different between the two regions, with cells in the spine having a significantly more heterogeneous distribution of actin accumulation outcomes (**Figure 3B, C**).

Most intriguing were the dynamics of N-cadherin localization, for which we observed two surprising patterns. First, in the anterior neural ectoderm we noticed that not only junctional *but also medial* N-cadherin dynamics strongly reflected those observed for actin (**Fig. 3D, D’, E, E’**). The latter result was surprising because classical cadherins such as N-cadherin are typically known for their function at apical cell-cell junctions. However, a new understanding of the cell-autonomous functions of cadherins, including N-cadherin, is now emerging (Ichikawa et al., 2020; Rebman et al., 2016; Sabatini et al., 2011). Indeed, so-called “extra-junctional cadherins” have been shown to interact with and even regulate the actomyosin cortex in cultured cells and *C. elegans* embryos (Padmanabhan et al., 2017; Sako et al., 1998; Wu et al., 2015). Our result suggests another noncanonical, non-junctional function for N-cadherin in the vertebrate neural plate.

In contrast to these patterns in the anterior neural ectoderm, N-cadherin dynamics in the posterior did *not* closely reflect actin dynamics. Though the pattern of actin accumulation in posterior neural ectoderm cells was more heterogeneous than that observed anteriorly, a majority of cells in the posterior plate accumulated both junctional and apical actin (**Fig. 3B, C**). However, neither junctional nor medial N-cadherin displayed significant accumulation in the posterior neural plate during the period of observation (**Fig. 3D, E**). These data provide a quantitative description of actin and N-cadherin dynamics in the developing neural plate and suggest that distinct mechanisms link actomyosin and cell adhesion to cell shape change in the anterior and posterior regions of the developing vertebrate neural ectoderm.

### Mosaic mutation of *shroom3* reveals distinct anterior and posterior phenotypes in the neural ectoderm

The differences in cell behaviors observed between anterior and posterior neural ectodermal regions mirrored the region-specific nature of NTDs in both humans and animal models. We therefore sought to understand how specific changes in cell behavior may explain region-specific NTDs. To this end, we examined the function of the *shroom3* gene, which is implicated in human NTDs and has variably penetrant effects on anterior and posterior neural tube closure, with more severe and penetrant anterior defects being observed in both mice and *Xenopus* (Deshwar et al., 2020; Haigo et al., 2003; Hildebrand and Soriano, 1999; Lemay et al., 2015).

F0 mutagenesis using CRISPR has recently emerged as a powerful tool in *Xenopus* and zebrafish, and mosaic crispants generated by targeted injections allow simultaneous observation of wild-type and crispant cellular phenotypes so that observations are automatically staged and synchronized (Aslan et al., 2017; Kakebeen et al., 2020; Kroll et al., 2021; Szenker-Ravi et al., 2018; Willsey et al., 2020). We therefore designed sgRNAs that effectively targeted the *shroom3* locus in *X. tropicalis* (**Methods Appendix Figure 1A, B**). Injection of these sgRNAs together with Cas9 protein into the 2 dorsal-animal blastomeres at the 8-cell stage induced neural tube closure defects (**Methods Appendix Figure 1C, D**), thus recapitulating knockdowns in *Xenopus* using MOs (Haigo et al., 2003), as well as the results in mouse genetic mutants (Hildebrand and Soriano, 1999). Critically, injections of sgRNA without Cas9 had no effect (**Methods Appendix Figure 1C, D**).

We therefore labeled the neural plate by injection of fluorescent reporters into both dorsal blastomeres at the 4-cell stage, and then injected a mixture of *shroom3*-targeted sgRNA, Cas9 protein, and membrane-BFP mRNA into one dorsal blastomere of 8-cell stage embryos to generate mosaic crispants (**Fig. 4A and see Methods Appendix Figure 1**), in which we identify *shroom3* crispant cells via membrane-BFP localization (**Fig. 4B**). Maps of the initial conformations of our imaged embryos and their *shroom3* crispant cell calls are located in **Methods Appendix Figure 2**. These maps allow us to systematically assess the relative locations of control and *shroom3* crispant cells along mosaic interfaces. For the purposes of brevity and simplicity, in this study we have not included in our analyses any cells or cell junctions situated along the mosaic interface (i.e. control cells that had *shroom3* crispant neighbors, *shroom3* crispant cells that had control neighbors, or junctions between control and *shroom3* crispant cells).

**Figure 4:**
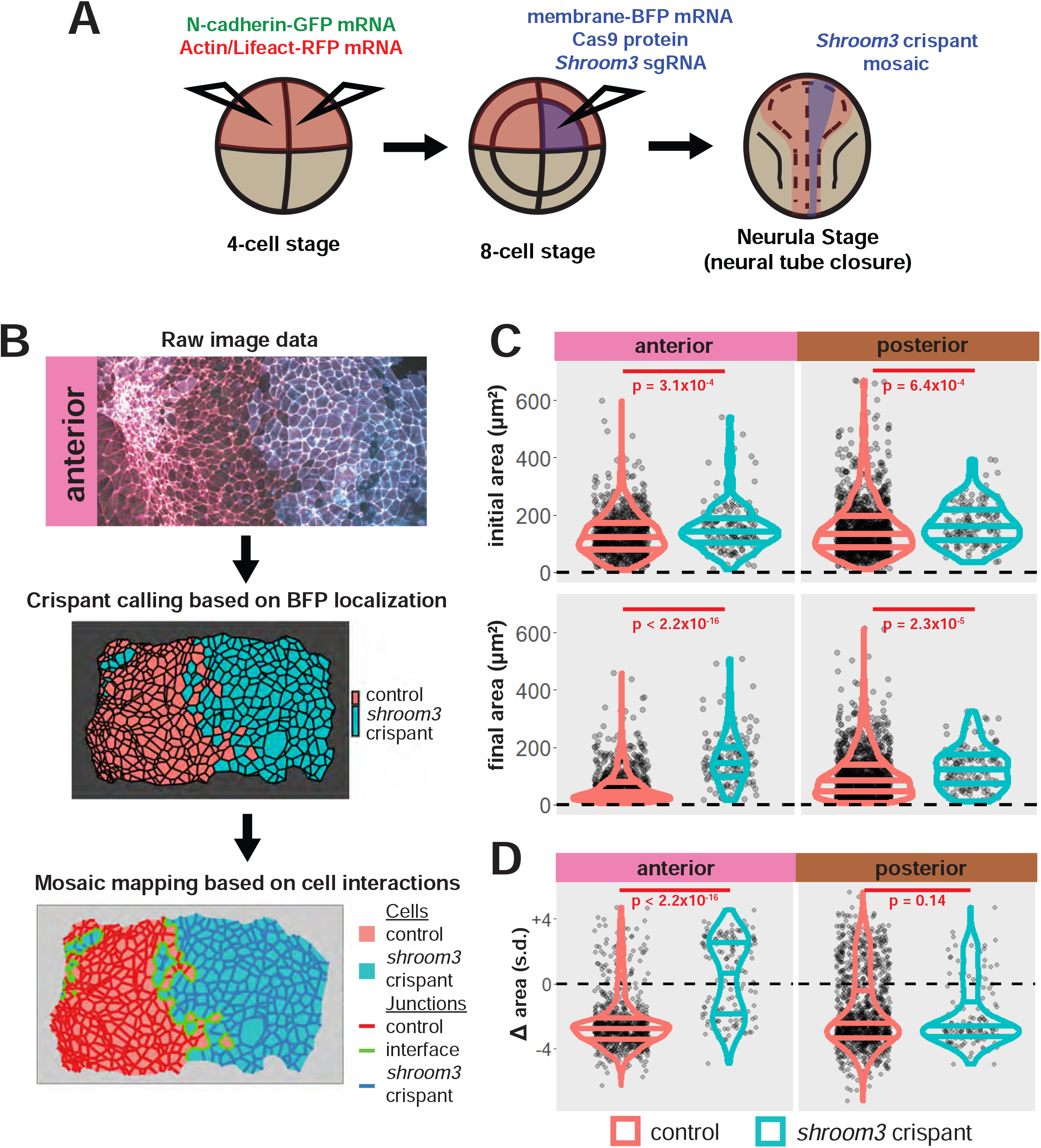
Disruption of *shroom3* via mosaic F0 CRISPR mosaic causes differential apical constriction phenotypes between regions of the neural ectoderm. **A**. Schematic of mosaic F0 CRISPR/Cas9 injections in *X. tropicalis* embryos. **B**. Workflow of identification and analysis of mosaic F0 crispants. **C**. Top row, distribution of initial area (square microns) of tracked cells from anterior (left) and posterior (right) embryos. Lower row, distribution of final area (square microns) of tracked cells. **D**. Distribution of overall change (Δ) in apical area (standardized) from all cells/embryos. In C & D, horizontal lines on density plots/violins indicate quartiles of distribution. Black circles are individual cells. Statistical comparisons performed by KS test. Cells situated along the mosaic interface were excluded from these analyses. s.d. = standard deviation.

Importantly, this F0 CRISPR-based approach recapitulated the known, specific cell biological phenotype of Shroom3 loss, as demonstrated previously in the neural tube of *Xenopus* morphants and mouse mutants (Lee et al., 2007; McGreevy et al., 2015; Plageman et al., 2010). Specifically, we observed a gross failure of anterior neural tube closure associated with defective apical constriction (**Fig. 4**). In the anterior region, *shroom3* crispant cells displayed significantly enlarged apical surfaces at the onset of our imaging (∼st. 13), and this phenotype grew more severe over time (**Fig. 4C**, left). Cells in the posterior region displayed a far more subtle phenotype, and while these cells were less constricted than controls at the onset of imaging, this phenotype did not become more severe over time (**Fig. 4C, right**). Thus, the change in apical area over the course of imaging was significantly disrupted in cells of the anterior neural plate, but surprisingly, was not significantly disrupted in the posterior neural plate during the period of imaging (**Fig. 4D**).

In summary, our F0 mosaic mutagenesis of *shroom3* elicited apical constriction phenotypes across the neural ectoderm, but the phenotype was far stronger in the anterior, consistent with the gross phenotype of highly penetrant anterior NTDs and far less penetrant posterior NTDs in both frog and mouse embryos lacking Shroom3 function (Haigo et al., 2003; Hildebrand and Soriano, 1999; McGreevy et al., 2015).

### Shroom3 links actin dynamics to effective apical constriction in the anterior neural ectoderm

A unified mechanism for apical constriction has emerged in recent years whereby apical constriction is driven by two discrete populations of actomyosin positioned either at apical cell-cell junctions or the medial apical cell surface (Martin and Goldstein, 2014). However, this model has been developed from studies in *C. elegans* and *Drosophila*, and the extent to which the model applies in vertebrates is unknown. We therefore examined our time-lapse data from the anterior neural ectoderm in order to understand the interplay of junctional and medial actin dynamics during vertebrate apical constriction.

First, we examined the gross change in medial and junctional actin over the period of imaging, roughly st. 13-18 (**Fig. 5A**). Wild-type cells tended to increase both medial and junctional actin localization over time, as expected (**Fig. 5B, C, red violin**). Surprisingly, despite the known role of Shroom3 in the control of actin accumulation (Haigo et al., 2003; Hildebrand, 2005), the majority of *shroom3* crispant cells actually *increased* both junctional and medial actin over the period of imaging (**Fig. 5B, C, blue violin**), though this increase was significantly reduced compared to controls (**Fig. 5B, C**).

**Figure 5:**
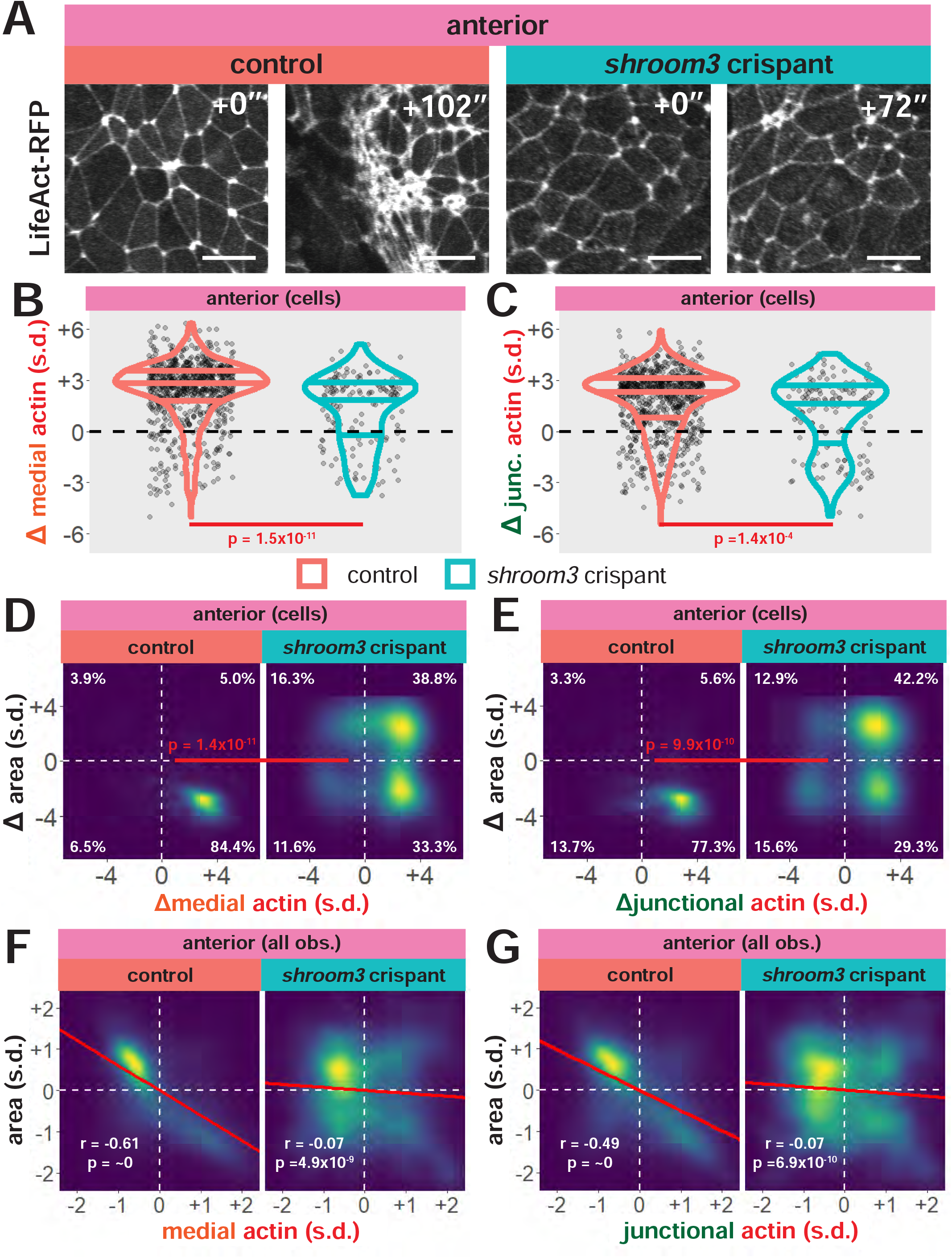
Medial actin accumulation drives apical constriction while loss of *shroom3* disrupts actin accumulation and constriction in the anterior neural ectoderm. **A**. Representative images of LifeAct/actin localization in control cells (left) and *shroom3* crispant cells (right) from the anterior region of the neural ectoderm. Scale bar = 15 microns. **B**. Distribution of overall change (Δ) in medial LifeAct/actin (standardized) from anterior cells. **C**. Distribution of overall change (Δ) in junctional LifeAct/actin (standardized) from anterior cells. In B & C, horizontal lines on density plots/violins indicate quartiles of distribution, black circles are individual cells, and statistical comparisons performed by KS test. **D & E**. 2D distribution of changes in apical area and medial (**D**) or junctional (**E**) LifeAct/actin (both standardized). Percentages in white indicate the percentage of total cells in each quadrant. Statistical comparisons performed by Peacock test, a 2D implementation of the KS test. **F & G**. 2D density plots of all observations of apical area versus medial (**F**) or junctional (**G**) LifeAct/actin for all cells within each group. Red lines indicate best-fit line through the observations. Statistics (r and p) are calculated for Pearson’s correlation. Cells situated along the mosaic interface were excluded from these analyses. s.d. = standard deviation.

To explore this surprising result in more detail, we directly compared changes in apical area with changes in actin intensity for each cell. In 838 control cells, we observed that the vast majority displayed a strong reduction in apical area and a strong increase in both junctional and medial actin intensity (**Fig. 5D, E**). As noted in the bulk statistics above, the majority of 147 *shroom3* crispant cells displayed increased actin intensity, but these crispant cells displayed a bimodal distribution of changes in apical area, with some cells constricting and other cells dilating (**Fig. 5D, E**). Thus, even cells that increase their apical area still also accumulate medial and junctional actin after loss of Shroom3.

Finally, for the most granular view, we plotted the correlation between standardized apical area and standardized actin intensity for all cells at all time points (N = ∼54k data points). For control cells, apical area displayed a very strong negative correlation with both medial and junctional actin intensity (**Fig. 5F, G**). Notably, however, medial actin was far more strongly correlated to apical area than was junctional actin, consistent with the key function of medial actin described in other systems. Interestingly, despite the relatively modest impact on overall actin accumulation after loss of Shroom3 (**Fig. 5B, C**), the normally strong correlations of apical surface area to medial and junctional actin dynamics were completely abolished in crispant cells (**Fig. 5F, G**). These results demonstrate that while Shroom3 may serve an overall role in actin assembly, this protein’s most critical function in neural tube closure *in vivo* is to convert apical actin dynamics into effective apical constriction.

### Loss of Shroom3 uncouples actin dynamics from N-cadherin in the anterior neural ectoderm

N-cadherin has been implicated in Shroom3 function and apical constriction in multiple contexts (Lang et al., 2014; Nandadasa et al., 2009; Plageman et al., 2011b), so we next asked how Shroom3 loss impacted N-cadherin dynamics during neural tube closure (**Fig 6A**). In contrast to the surprisingly modest change observed in actin intensity (**Fig. 5B,C**), bulk measurements revealed that *shroom3* crispant cells displayed a profound failure to accumulate medial N-cadherin, and in fact the majority of cells actually reduce N-cadherin levels (**Fig. 6B, blue violin**). However, junctional N-cadherin dynamics were more modestly disrupted by Shroom3 loss, with many *shroom3* crispant cells showing some degree of junctional N-cadherin accumulation (**Fig. 6C, blue violin**).

**Figure 6:**
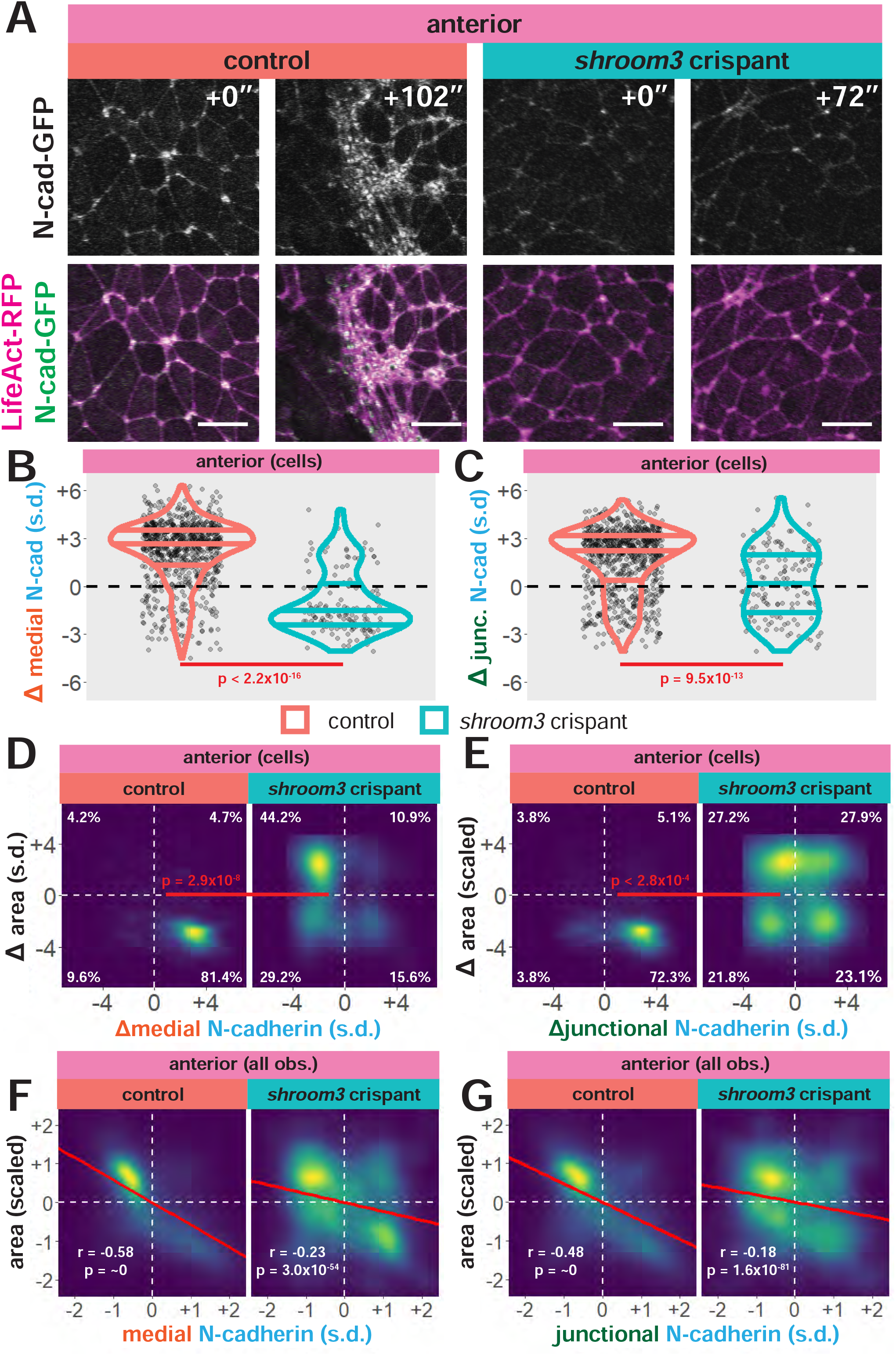
Medial N-cadherin accumulation is severely disrupted in anterior *shroom3* crispant cells that fail to apically constrict. **A**. Representative images of N-cadherin localization in control cells (left) and *shroom3* crispant cells (right) from the anterior region of the neural ectoderm. Scale bar = 15 microns. **B**. Distribution of overall change (Δ) in medial N-cadherin (standardized) from anterior cells. **C**. Distribution of overall change (Δ) in junctional N-cadherin (standardized) from anterior cells. In B & C, horizontal lines on density plots/violins indicate quartiles of distribution, black circles are individual cells, and statistical comparisons performed by KS test. **D & E**, 2D distribution of changes in apical area and medial (**D**) or junctional (**E**) N-cadherin (both standardized). Percentages in white indicate the percentage of total cells in each quadrant. Statistical comparisons performed by Peacock test, a 2D implementation of the KS test. **F & G**. 2D density plots of all observations of apical area versus medial (**F**) or junctional (**G**) N-cadherin for all cells within each group. Red lines indicate best-fit line through the observations. Statistics (r and p) are calculated for Pearson’s correlation. Cells situated along the mosaic interface were excluded from these analyses. s.d. = standard deviation.

Next, we performed direct comparisons of changes in apical area to changes in N-cadherin intensity. The vast majority of control cells (81%) reduced their apical area while increasing medial N-cadherin localization (**Fig. 6D, orange**), while *shroom3* crispant cells displayed the converse behavior, with a plurality of cells displaying apical surface dilation over time coupled to a strong *reduction* in medial N-cadherin intensity over time (**Fig. 6D, blue**). Consistent with the bulk statistics above, junctional N-cadherin displayed a distinct phenotype, with cells displaying all possible combinations of apical surface behavior and N-cadherin intensity profiles. (**Fig. 6E**). Finally, correlations of apical area to N-cadherin intensity for all cells at all time-points revealed very strong correlations for both medial and junctional N-cadherin in controls, and these correlations were significantly disrupted by Shroom3 loss (**Fig. 6F, G**).

Notably, in all of our analyses, we observed that medial N-cadherin intensity was more strongly affected than was junctional N-cadherin, medial actin or junctional actin, which is intriguing in light of recent reports of key roles for extra-junctional cadherins in actin-dependent processes (Padmanabhan et al., 2017; Sabatini et al., 2011). To further explore this issue, we compared actin and N-cadherin dynamics directly (N = ∼54k observations). In control cells, both medial and junctional N-cadherin dynamics display very robust correlation to actin (**Fig. 7**), consistent with our data above. Importantly, however, after Shroom3 loss this correlation was severely disrupted for the medial populations, while the disruptive effect on junctional correlations was far less robust (**Fig. 7**).

**Figure 7:**
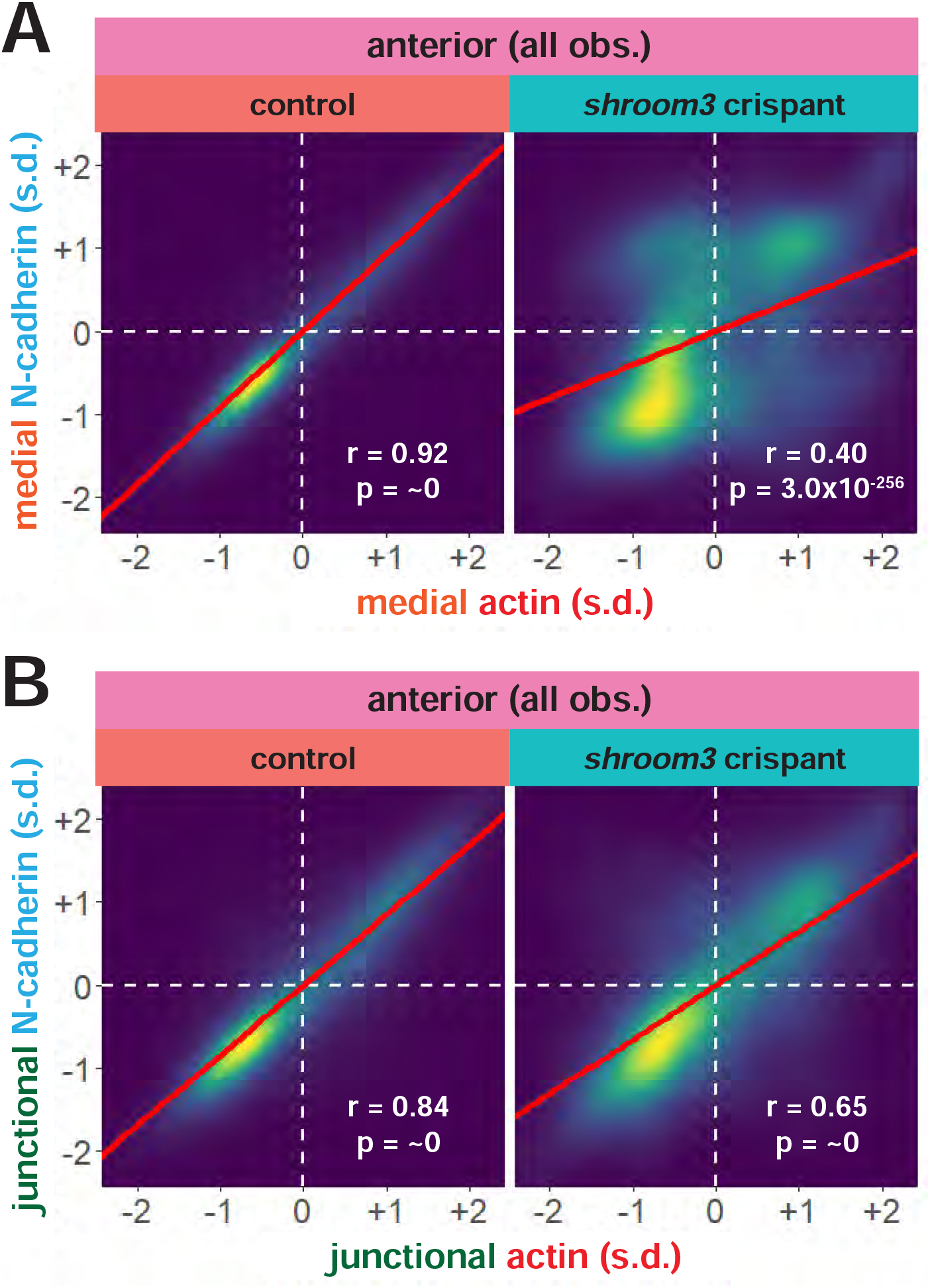
Actin and N-cadherin accumulation are uncoupled in anterior *shroom3* crispant cells. **A & B**. 2D density plots of all observations of medial (**A**) or junctional (**B**) LifeAct/actin versus N-cadherin for all cells within each group. Red lines indicate best-fit line through the observations. Statistics (r and p) are calculated for Pearson’s correlation. Cells situated along the mosaic interface were excluded from these analyses. s.d. = standard deviation.

Together, these results suggest that the primary effect of *shroom3* loss in the anterior neural plate is not on actomyosin contractility *per se*, but rather on the coupling of medial actomyosin contractility with medial N-cadherin accumulation. This result further highlights the possibility of an extra-junctional role for N-cadherin in apical constriction and moreover provides a mechanism to explain our findings above (**Fig. 5D**) that Shroom3 links actin dynamics to changes in apical area.

### In the posterior neural ectoderm, Shroom3 controls actin and N-cadherin, but not apical constriction

As noted above, *shroom3* crispant cells in the posterior neural ectoderm did not display appreciable defects in the change in apical area over the period of our imaging, though cells were overall less constricted than in controls (**Fig. 4C, D**). We reasoned that this result may reflect the complex interplay of Shroom3-mediated apical constriction and PCP-dependent convergent extension cell behaviors (e.g. (McGreevy et al., 2015; Nishimura et al., 2012)). We therefore explored our image data for new insights into the role of Shroom3 in the posterior neural ectoderm.

Both junctional and medial actin accumulated over the course of our imaging in control cells in the posterior neural ectoderm, as expected (**Fig. 8A-C, red violins**). In addition, comparisons of changes in area to changes in actin localization revealed that the vast majority of cells decreased their apical area and increased both medial and junctional actin localization (**Fig. 8D, E**). Interestingly, however, when comparing the relationship between apical area and actin across all time points in posterior control cells, we found similarly robust correlations for both medial and junctional actin (**Fig. 8F, G**). These correlations are in contrast with our data for the anterior neural ectoderm, where medial actin is more strongly correlated with apical area than junctional actin in control cells (see **Fig. 5F, G**).

**Figure 8:**
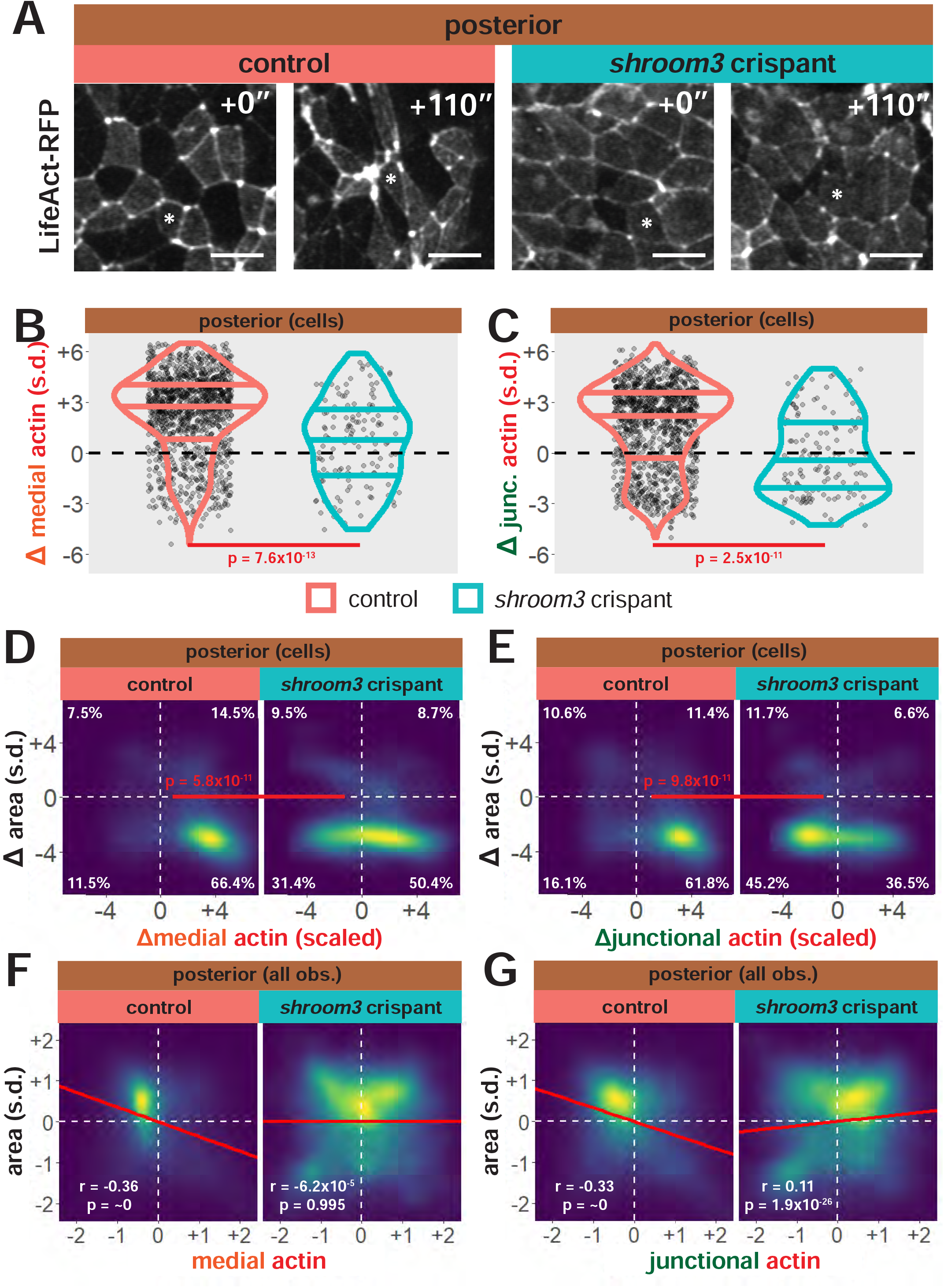
Loss of *shroom3* disrupts actin dynamics in the posterior neural ectoderm. **A**. Representative images of LifeAct/actin localization in control cells (left) and *shroom3* crispant cells (right) from the posterior region of the neural ectoderm. White asterisks mark the same cell in each embryo. Scale bar = 15 microns. **B**. Distribution of overall change (Δ) in medial LifeAct/actin (standardized) from posterior cells. **C**. Distribution of overall change (Δ) in junctional LifeAct/actin (standardized) from anterior cells. In B & C, horizontal lines on density plots/violins indicate quartiles of distribution, black circles are individual cells, and statistical comparisons performed by KS test. **D & E**. 2D distribution of changes in apical area and medial (**D**) or junctional (**E**) LifeAct/actin (both standardized). Percentages in white indicate the percentage of total cells in each quadrant. Statistical comparisons performed by Peacock test, a 2D implementation of the KS test. **F & G**. 2D density plots of all observations of apical area versus medial (**F**) or junctional (**G**) LifeAct/actin for all cells within each group. Red lines indicate best-fit line through the observations. Statistics (r and p) are calculated for Pearson’s correlation. Cells situated along the mosaic interface were excluded from these analyses. s.d. = standard deviation.

Examination of actin dynamics in *shroom3* crispant cells revealed clear differences between anterior and posterior neuroepithelial cells. Most strikingly, despite the far less robust apical constriction defect in posterior *shroom3* crispant cells, we observed a much *stronger* defect in accumulation of both medial and junctional actin (compare **Fig. 8B, C** with **Fig. 5B, C**). Moreover, comparing changes in actin to changes in area revealed that many *shroom3* crispant cells undergoing robust apical constriction in the posterior neural plate nonetheless displayed an obvious *loss* of apical actin, suggesting that overall changes in actin localization do not drive apical constriction in posterior *shroom3* crispant cells (**Fig. 8D,E**). In agreement with this idea, both medial and junctional actin localization are relatively poorly correlated with apical area in posterior *shroom3* crispant cells (**Fig. 8F, G**).

The dynamics of N-cadherin localization revealed an even more pronounced difference between posterior and anterior (**Fig. 9A**). In control embryos, N-cadherin in the posterior neural ectoderm displayed a far less robust pattern of change over the course of imaging (**Fig. 9B, C, red violins**), a clear departure from what was observed anteriorly (Compare to **Fig. 6B, C**). In fact, direct comparisons of overall change in area to overall changes in N-cadherin localization in the posterior revealed two distinct populations of control cells, one in which cells decrease apical area and increased N-cadherin, and another which decreased apical area but also decreased N-cadherin **(Fig. 9D**). A similar pattern was observed for junctional N-cadherin (**Fig. 9E**). Accordingly, comparisons of apical area to medial or junction N-cadherin across all time points revealed essentially no correlations (**Fig. 9F, G**).

**Figure 9:**
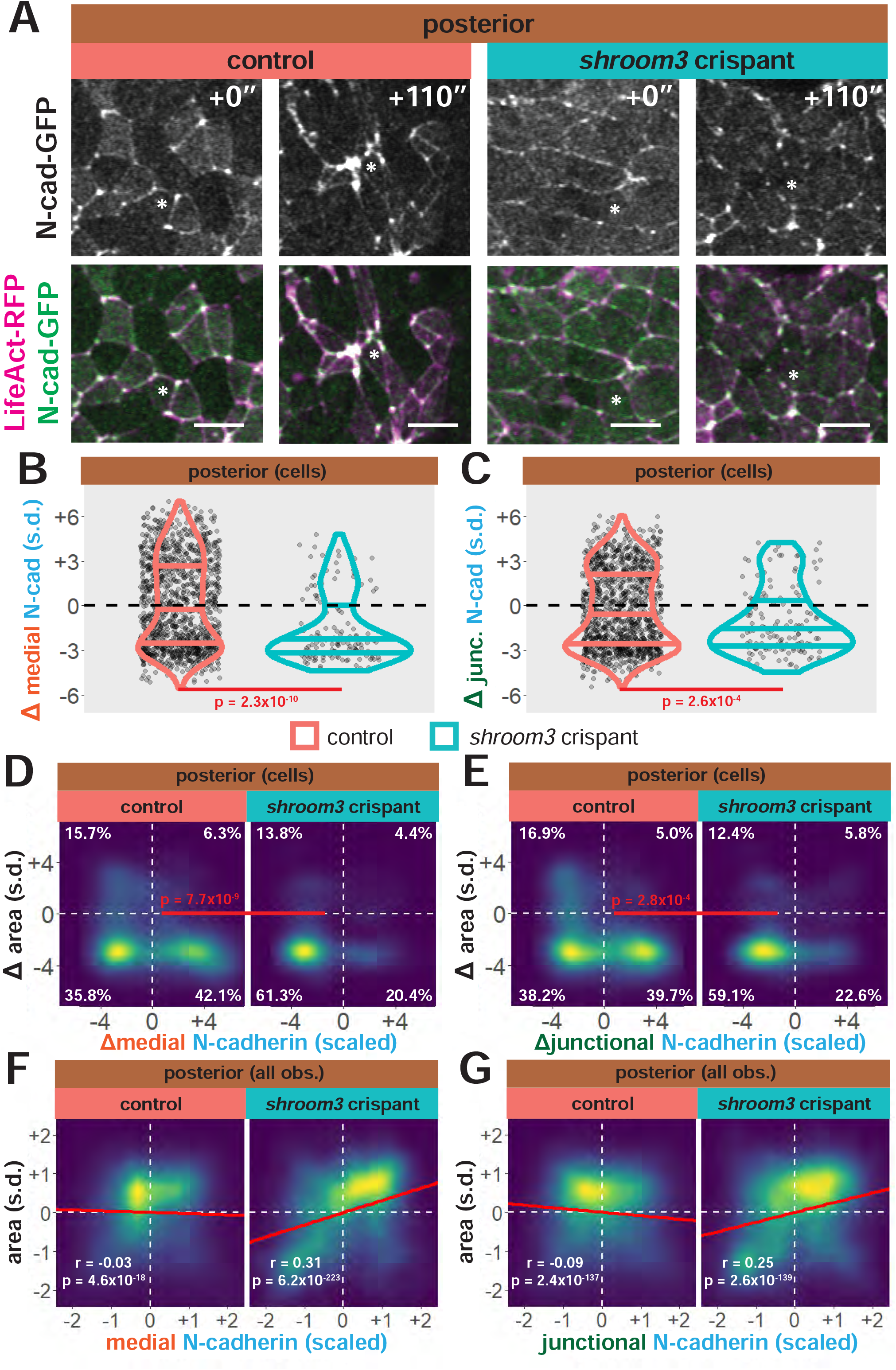
N-cadherin dynamics are highly heterogenous in the posterior neural ectoderm and poorly correlated with apical constriction. **A**. Representative images of N-cadherin localization in control cells (left) and *shroom3* crispant cells (right) from the posterior region of the neural ectoderm. White asterisks mark the same cell in each embryo. Scale bar = 15 microns. **B**. Distribution of overall change (Δ) in medial N-cadherin (standardized) from posterior cells. C, distribution of overall change (Δ) in junctional N-cadherin (standardized) from posterior cells. In B & C, horizontal lines on density plots/violins indicate quartiles of distribution, black circles are individual cells, and statistical comparisons performed by KS test. **D & E**. 2D distribution of changes in apical area and medial (**D**) or junctional (**E**) N-cadherin (both standardized). Percentages in white indicate the percentage of total cells in each quadrant. Statistical comparisons performed by Peacock test, a 2D implementation of the KS test. **F & G**. 2D density plots of all observations of apical area versus medial (**F**) or junctional (**G**) N-cadherin for all cells within each group. Red lines indicate best-fit line through the observations. Statistics (r and p) are calculated for Pearson’s correlation. Cells situated along the mosaic interface were excluded from these analyses. s.d. = standard deviation.

Interestingly, *shroom3* crispant cells in the posterior neural plate displayed a fairly uniform behavior, constricting normally, but with nearly all cells also reducing both medial and junctional N-cadherin intensity (**Fig. 9B, C, blue violins, D, E**). Thus, the impact of Shroom3 loss on N-cadherin dynamics is stronger in the posterior than the anterior, even though the effect on apical constriction is far weaker. Together, these results suggest that while a fairly direct relationship between Shroom3, actin, and N-cadherin governs apical constriction anteriorly, the situation is far more complex in the posterior neural ectoderm. One possible explanation for this added complexity is that PCP proteins, such as Vangl2, are known to have a stronger loss-of-function phenotypes in the posterior and likely exert stronger control over cell-cell junction behavior than Shroom3 in posterior neural tube closure; loss of Shroom3 may thus result in aberrant actomyosin localization but relatively unperturbed apical constriction.

### Cell-cell junction behaviors display distinct differences between the anterior and posterior regions of the neural ectoderm

While medial actomyosin appears to be a stronger driver of apical constriction in the anterior neural ectoderm than junctional actomyosin, the primary site of actomyosin activity in the posterior neural ectoderm was unclear. Both actin and N-cadherin accumulation were highly variable at both the medial and junctional domains of posterior cells, and actin and N-cadherin localization were relatively poorly correlated with apical area (**Figs. 8, 9**). To gain additional insights into posterior neural cell behaviors, we used our dataset to quantify the specific behaviors of each individual cell-cell junction in our dataset, incorporating nearly half a million observations from over 10,000 junctions.

At the bulk level, junctions in the anterior on average decreased in length, as expected and this trend was severely disrupted in *shroom3* crispant junctions (p<2.2×10^−16^)(**Fig. 10A**). In the posterior, however, we observed roughly equal growth and shrinkage of control junctions (**Fig. 10A**). In contrast to anterior *shroom3* junctions, posterior *shroom3* crispant junctions did not show a change in overall junction shrinkage and growth relative to control junctions (p=0.2121)(**Fig 10A**).

**Figure 10:**
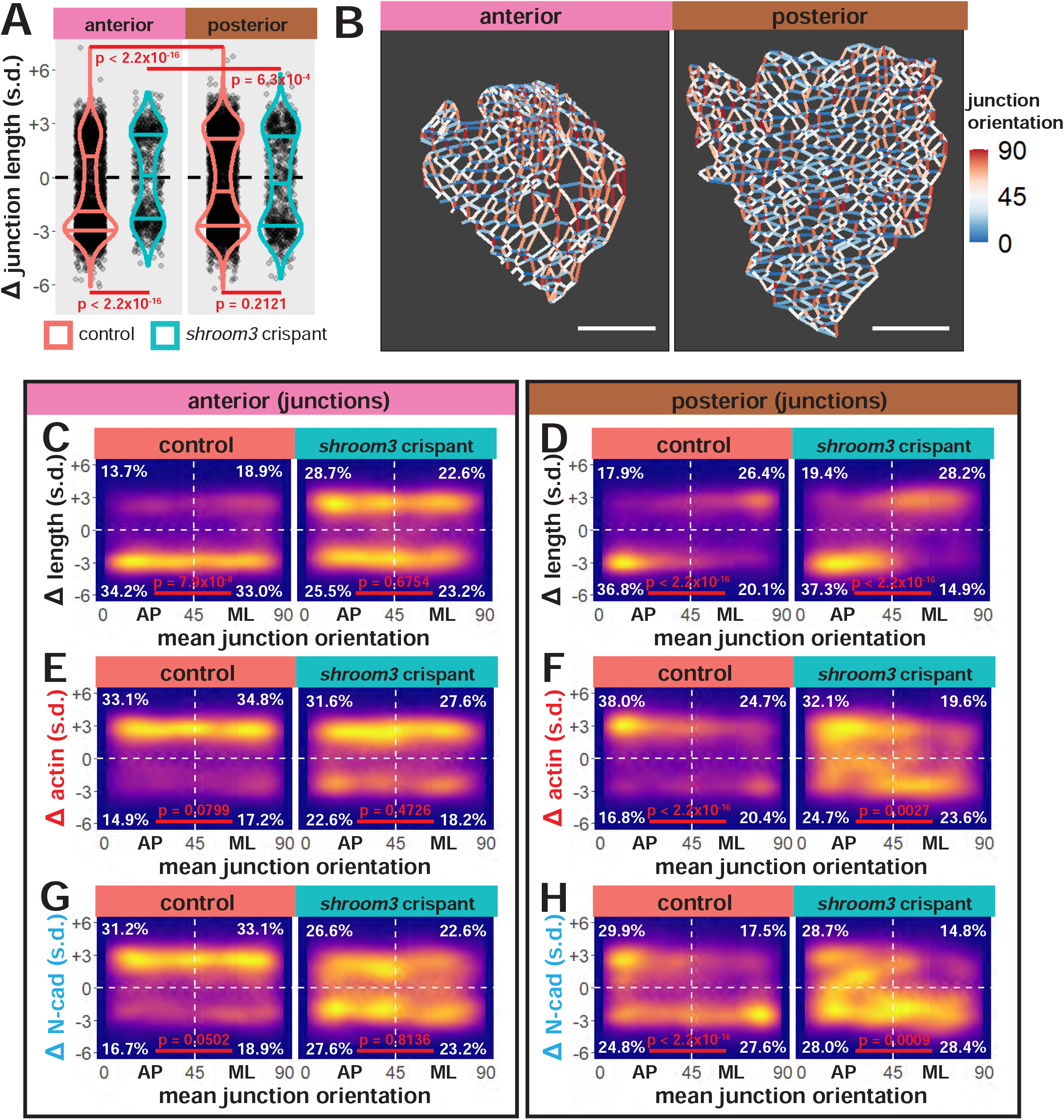
Individual junction behaviors are more strongly polarized in the posterior neural ectoderm. **A**. Distribution of overall change (Δ) in junction length (standardized) from the anterior (left) and posterior (right) regions. Horizontal lines on density plots/violins indicate quartiles of distribution, black circles are individual cells, and statistical comparisons performed by KS test. **B**. Junction orientation from initial frames of control embryos from Figure 2. Scale bars = 100 microns. **C & D**. 2D density plots of all observations of mean junction orientation over time versus overall change (Δ) in junction length (standardized) for all junctions within each group. **E & F**. 2D density plots of all observations of mean junction orientation over time versus overall change (Δ) in junction actin (standardized) for all junctions within each group. **G & H**. 2D density plots of all observations of mean junction orientation over time versus overall change (Δ) in junction N-cadherin (standardized) for all junctions within each group. Percentages in white indicate the percentage of total cells in each quadrant. Statistical comparisons performed by Peacock test, a 2D implementation of the KS test. s.d. = standard deviation.

Junction behaviors during epithelial morphogenesis are frequently polarized with respect to the embryonic axes (Pinheiro and Bellaïche, 2018), so we assigned orientations to all junctions in our dataset. Junctions with a mean orientation less than 45° were designated anteroposterior (AP), as they represent the anterior face of one cell abutting the posterior face of a neighboring cell; junctions with mean orientations greater than 45° were designated as mediolateral (ML)(**Fig. 10B**).

In the anterior neural plate, the majority of control junctions displayed robust shrinkage with minor polarization, as evidenced by a difference in distribution of shrinkage and growth between AP and ML junctions (**Fig. 10C**). More specifically, slightly more ML junctions grew than AP junctions, indicating that some degree of tissue elongation or isotropic apical constriction occurs in the anterior neural ectoderm. By contrast, anterior *shroom3* crispant junctions showed no clear bias towards shrinkage or growth in either AP or ML junctions and no polarization of AP versus ML junctions (**Fig. 10C**).

While changes in junction length were somewhat polarized in anterior control junctions, neither actin (**Fig. 10E**) or N-cadherin (**Fig. 10G**) dynamics were significantly polarized in those same junctions. Anterior control junctions accumulated both actin and N-cadherin at a similar frequency to which they shrank. Actin and N-cadherin dynamics in anterior *shroom3* crispant junctions were not polarized, similar to control junctions (**Fig. 10E, G**). However, the proportion of junctions that reduced actin and N-cadherin localization decreased in crispant junctions, similar to the increased proportion of crispant junctions that grew over time (**Fig. 10C**). Overall, these results again suggest that Shroom3 may control actomyosin and N-cadherin accumulation at anterior cell-cell junctions.

In the posterior neural plate, we observed far more complex patterns of junction behavior. First, junction shrinkage in the posterior neural plate was highly polarized, with AP junctions predominantly shortening and ML junctions predominantly growing (**Fig. 10D**). These patterns of junction behavior are hallmarks of convergent extension and are consistent with previous smaller scale studies of these behaviors in the neural ectoderm (Butler and Wallingford, 2018b; Nishimura et al., 2012; Williams et al., 2014a) and in other tissues (Bertet et al., 2004; Blankenship et al., 2006; Lienkamp et al., 2012). Strikingly, we observed a very similar pattern of behavior in posterior *shroom3* junctions, indicating that Shroom3 has little to no role in polarizing junction behaviors in the posterior neural ectoderm (**Fig. 10D**).

However, examination of the patterns of actin and N-cadherin dynamics in the posterior neural ectoderm revealed a more complex phenotype. While we still observed polarization of actin accumulation in AP versus ML control junctions, actin tended to accumulate in both AP and ML junctions (**Fig. 10F**) despite the fact that ML junctions tended to grow (**Fig. 10F**). Strikingly, despite the fact that *shroom3* crispant junctions changed their length in a highly polarized manner (**Fig. 10D**), actin accumulation was depolarized to a significant degree in crispant junctions (**Fig. 10F**), consistent with data from still images in the neural plate of mouse *shroom3* mutants (McGreevy et al., 2015). This defect seems mainly to be due to an increase in actin clearance in *shroom3* AP junctions, despite our observation that *shroom3* AP junctions shrink to a very similar degree as control AP junctions (**Fig. 10D**). These results suggest a disconnect between actin accumulation and junction shrinkage in the posterior neural ectoderm that is amplified upon loss of Shroom3.

Even more interesting were the patterns of N-cadherin dynamics. Recall that in cell-level data, we observed nearly equal populations of cells that gained and lost junctional N-cadherin over time (**Fig. 9E**), which we anticipated might reflect junction polarity. What we discovered was that N-cadherin accumulation was again polarized in AP versus ML control junctions, such that shortening AP-oriented junctions were equally likely to accumulate N-cadherin, while ML junctions, which preferentially grow, were far more likely to clear N-cadherin (**Fig. 10H**). Again, *shroom3* junctions presented an unusual phenotype; N-cadherin dynamics were less polarized than length changes in posterior crispant junctions, and the defect again seemed mainly to be an increase in N-cadherin *clearance* at *shroom3* AP junctions (**Fig. 10H**). Thus, posterior *shroom3* crispant junctions display largely normal polarized changes in length (**Fig. 10D**) but show reduced or altered polarity for both actin and N-cadherin dynamics (**Fig. 10F, H**).

Together, these results again paint a vivid picture of differences between cell behaviors in the anterior and posterior neural ectoderm. Both control and *shroom3* junctions in the anterior behaved mostly as expected; the majority of both AP and ML control junctions shrank over time, and loss of Shroom3 disrupted junction shrinkage as well as actin and N-cadherin dynamics in a predictable pattern based on whole-cell data. In contrast, shrinkage and growth were highly polarized in the posterior neural ectoderm in both control and *shroom3* crispant junctions, despite polarization of actin and N-cadherin dynamics being reduced in crispant junctions.

## Conclusion

Understanding the cellular mechanisms contributing to neural tube closure has long challenged embryologists due to the relatively large number of cells involved and the heterogeneity of their behaviors. Indeed, some of the earliest uses of computer simulation in developmental biology focused on understanding the degree to which neural ectoderm cells constrict their apical surfaces during neural tube closure (Jacobson and Gordon, 1976). Since then, our understanding of the genetics and cellular mechanisms of neural tube closure has broadly improved. Hundreds of mutations affecting neural tube closure have been identified (Harris and Juriloff, 2010), and interactions between neural tube closure genes are becoming better understood (McGreevy et al., 2015; Murdoch et al., 2014). However, while time lapse imaging of single cell behaviors in chicks, frogs, and mice have provides several key insights (Butler and Wallingford, 2018a; Christodoulou and Skourides, 2015; Davidson and Keller, 1999; Galea et al., 2017; Massarwa and Niswander, 2013; Molè et al., 2020; Ossipova et al., 2015; Pyrgaki et al., 2010; Wallingford and Harland, 2002; Williams et al., 2014b), our understanding of the cell biology of neural tube closure still lags substantial behind our understanding of the genetics.

Recent research has begun to bridge the gap between the breadth of our genetic resources and the scale of neural tube closure as a process; *in toto* imaging has captured neural tube closure as one facet of overall mouse development (McDole et al., 2018), tissue-scale imaging of has revealed the cell biological basis of ciliopathy-related NTDs (Brooks et al., 2020). In a complement to these advancements, we have used high-resolution, but tissue-scale, time-lapse imaging of both actin and N-cadherin followed by cell-tracking analysis to gain several new mechanistic insights into the process of apical constriction during neural tube closure.

First, we show that anterior and posterior neural ectoderm cells undergo apical constriction to differing degrees and by different constrictive mechanisms. Further, we show that mosaic mutation of *shroom3* similarly results in different phenotypes in the anterior and posterior. This is consistent with how disruption of *shroom3* has been shown to have stronger effects on anterior neural tube closure (Hildebrand 1999, Haigo 2003, McGreevy 2015). Conversely, disruption of PCP signaling has been shown to have stronger defects in posterior neural tube closure (e.g. (Galea et al., 2018; Murdoch et al., 2014)). Applying the approach described here to analysis of PCP protein localization should be highly illuminating.

Second, we show that changes in apical area in the anterior neural ectoderm are more strongly correlated with changes in medial actomyosin localization versus junctional actomyosin, indicating that contractility driving apical constriction is more likely to be generated at the medial cell surface. Our results on medial actin localization are largely consistent with smaller scale studies in both *Xenopus* and mice (Christodolou 2015, Suzuki 2017, Brown 2018) and moreover reflect mechanisms described in more detail in the context of *Drosophila* and *C. elegans* gastrulation (Martin et al., 2009; Roh-Johnson et al., 2012). Conversely, our data shows that both medial and junctional actin localization have similar correlations with apical area in the posterior neural ectoderm, suggesting actomyosin contractility may be more balanced across these medial and junctional domains in this region.

Third, we show that N-cadherin accumulates at the medial surfaces of constricting cells in the anterior neural ectoderm, but that N-cadherin localization poorly correlates with apical constriction in the posterior. Interestingly, our mechanistic analysis suggests that anterior *shroom3* crispant cells fail to constrict less from a lack of actomyosin accumulation and more from an inability to accumulate N-cadherin at the medial surface of cells. This is consistent with a previously reported genetic interaction between *shroom3* and *N-cadherin* (Plageman et al., 2011b), and suggests that a Shroom3-N-cadherin pathway may drive anterior apical constriction during neural tube closure.

The study of developmental biology in the 21^st^ century is marked by explosive increases in the size of experiments – advancements in proteomics and transcriptomics are providing an unprecedented depth to our understanding of the molecular workings of cells in embryos. Thus, it is of special importance that we use advances in imaging and data analysis to ask what those cells *actually do and how they do it*. The work here is therefore significant for providing quantitative insights into the interplay of gene function, protein localization and cell behavior during a biomedically important process in vertebrate embryogenesis.

## Acknowledgments

Special thanks to Pavak Shah and Claire McWhite for assistance with coding and data analysis and to the Wallingford lab for manuscript comments. This work was funded by NICHD Ruth L. Kirschstein NRSA F32 HD094521 for AB and R01HD099191 to JW.

## Materials and Methods

### Animals

Wild-type *Xenopus tropicalis* frogs were obtained from the National Xenopus Resource, Woods Hole, MA (Horb et al., 2019).

### Injections

Wild-type *X. tropicalis* eggs were fertilized *in vitro* using sperm from wild-type *X. tropicalis* males using standard methods.

*X. tropicalis* embryos were moved to 1/9x MMR + 2% Ficoll, then injected in both dorsal blastomeres at the 4-cell stage with 50pg LifeAct-RFP mRNA and 150pg *Xenopus* N-cadherin-GFP mRNA. In CRISPR-injected embryos, at the 8-cell stage, one dorsal blastomere was injected with 1ng Cas9 protein (PNA Bio), 250pg *shroom3*-targeted sgRNA (Synthego)(**Methods Appendix Fig. 1A**), and 60pg membrane(CAAX)-BFP mRNA.

### CRISPR genotyping

To test the efficacy of our CRISPR injections *in vivo*, we injected wild-type *X. tropicalis* with the above-described Cas9+sgRNA combination in the following cells and stages: 2x injections into 1-cell stage embryos, 1x injections into each blastomere of 2-cell stage embryos, 1x injections into 2 blastomeres of the 4-cell stage embryo, and 1x injections into all blastomeres of 4-cell stage embryos (**Methods Appendix Fig. 1B**). Uninjected embryos that did not receive any Cas9+sgRNA were used as controls.

Embryos were allowed to grow to approximately Nieuwkoop and Faber (NF) stage 40 and then were subjected to whole-embryo DNA extraction. PCR products spanning the *shroom3* target site were generated from each embryo and separated by capillary electrophoresis for fragment analysis. Fragment analysis data was analyzed in R using the *Fragman* package (Covarrubias-Pazaran et al., 2016).

Uninjected embryos did not have any indel products at the *shroom3* locus and thus produced one sharp peak at 431 base pairs, corresponding to the size of the wild-type *shroom3* PCR product (**Methods Appendix Fig. 1B**). By contrast, CRISPR-injected embryos returned little to no PCR products at this size (**Methods Appendix Fig. 1B**, dashed red line), indicating that the *shroom3* target site was being efficiently cut by Cas9 and repaired by error-prone pathways.

The exception to this were the embryos of which only 2 blastomeres at the 4-cell stage were injected with Cas9+sgRNA; as expected, fragments of the wild-type size were detected that theoretically correspond to the uninjected blastomere lineages (**Methods Appendix Fig. 1B**). Overall, these results indicate that our Cas9+sgRNA combination efficiently cleaves the *shroom3* target site *in vivo*.

### Imaging

Injected embryos were held at 25C until they reached NF stage 12.5. At NF stage 12.5, vitelline envelopes were removed from embryos and embryos were allowed to “relax” for 30 minutes.

Embryos were then mounted in imaging chambers and positioned for imaging of either the anterior or posterior neural plate.

Embryos were imaged on a Nikon A1R confocal microscope using the resonant scanner mode. Image quality, Z-stacking, and XY tiling were balanced to generate optimal 3D images of the neural plate at a rate of 1 frame per minute. Ultimately, movies of 7 embryos were of sufficient length and quality for analysis. Tissue geometry and *shroom3* crispant calling of the initial frame of each of these embryos is presented in **Methods Appendix Fig. 2**.

### Image Analysis

Raw 3D images were projected to 2D via maximum intensity and underwent initial segmentation of cell boundaries using the FIJI plugin Tissue Analyzer (Aigouy et al., 2010; Aigouy et al., 2016). The segmentation of an initial frame was hand-corrected, and this hand-corrected segmentation was used to train a classifier using the program CSML (Ota et al., 2018). CSML was used to generate segmentation for subsequent frames, which were then further hand-corrected in Tissue Analyzer.

After hand-correction, Tissue Analyzer was used to track both cell surfaces and cell junctions, then generate a database of measurements of size and fluorescent intensities for each cell and junction over time. Values for medial and junctional localization of imaged markers in cells were calculated as average pixel fluorescence intensity across the entirety of each respective domain. Similarly, localization of imaged markers to individual junctions was calculated as an average across the entire junction.

For individual junctions, errors in junction length caused by Z-displacement and projection were corrected in Matlab (code courtesy of Pavak Shah).

Tissue Analyzer databases were imported to R and further analyzed and manipulated primarily using the *tidyverse* package (R Core Team, 2020; Wickham et al., 2019).

### Data Analysis

Cell tracks shorter than 30 frames and junction tracks shorter than 15 frames were discarded.

Embryo “b” had a fluorescence anomaly during imaging that resulted in a reduction in overall fluorescence followed by a recovery (**Methods Appendix Fig. 3A,B**). Cells were tracked through the anomaly (**Methods Appendix Fig. 3C**), but fluorescent values for the frames 23 to 45 were discarded (**Methods Appendix Fig. 3**, red dashed box).

Cells were determined to be control versus *shroom3* crispant by a membrane-BFP localization threshold specific to each embryo (**Fig. 4B**, middle panel). Crispant calls were then manually annotated in cases along the mosaic interface where thresholding produced crispant calls deemed incorrect.

Individual junctions were determined to be control versus *shroom3* crispant versus mosaic interface based on the status of the cells the junction was situated between. Control junctions are situated between two control cells, *shroom3* crispant junctions are situated between two *shroom3* crispant cells, and mosaic interface junctions are situated between a control and a *shroom3* crispant cell (**Fig. 4B**, lower panel). In this study, we excluded cells and junctions at the mosaic interface from our analysis.

Junction orientations were corrected so that the mediolateral axis of the embryo was set at 0° and the anteroposterior axis of the embryo was set at 90° (**Figure 10A**).

Individual cell and junction tracks were smoothed by averaging over a 7 frame/minute window (**Methods Appendix Fig. 4B**). Individual cell tracks were further mean-centered and standardized so that variables are measured in standard deviations rather than fluorescence or size units (**Methods Appendix Fig. 4C**). This standardization allows us to analyze dynamics of cell size and protein localization across a population of cells while controlling for initial size and fluorescence of cells. In an example embryo, cells begin and end tracking with a variety of apical surface areas (**Methods Appendix Fig. 4D’**), but once the cell tracks are mean-centered and standardized it becomes clear that the cells are behaving similarly at a population level (**Methods Appendix Fig. 4C”**).

1D statistical comparisons were performed using the Kolmogorov-Smirnov (K-S) test via the R *ks*.*test* function (R Core Team, 2020). 2D statistical comparisons were performed using the Peacock Test, a 2D implementation of the K-S test (Fasano and Franceschini, 1987; Peacock, 1983). Peacock tests were performed using the R *Peacock*.*test* package (Xiao, 2016; Xiao, 2017).

---

**Methods Appendix Figure 1:**
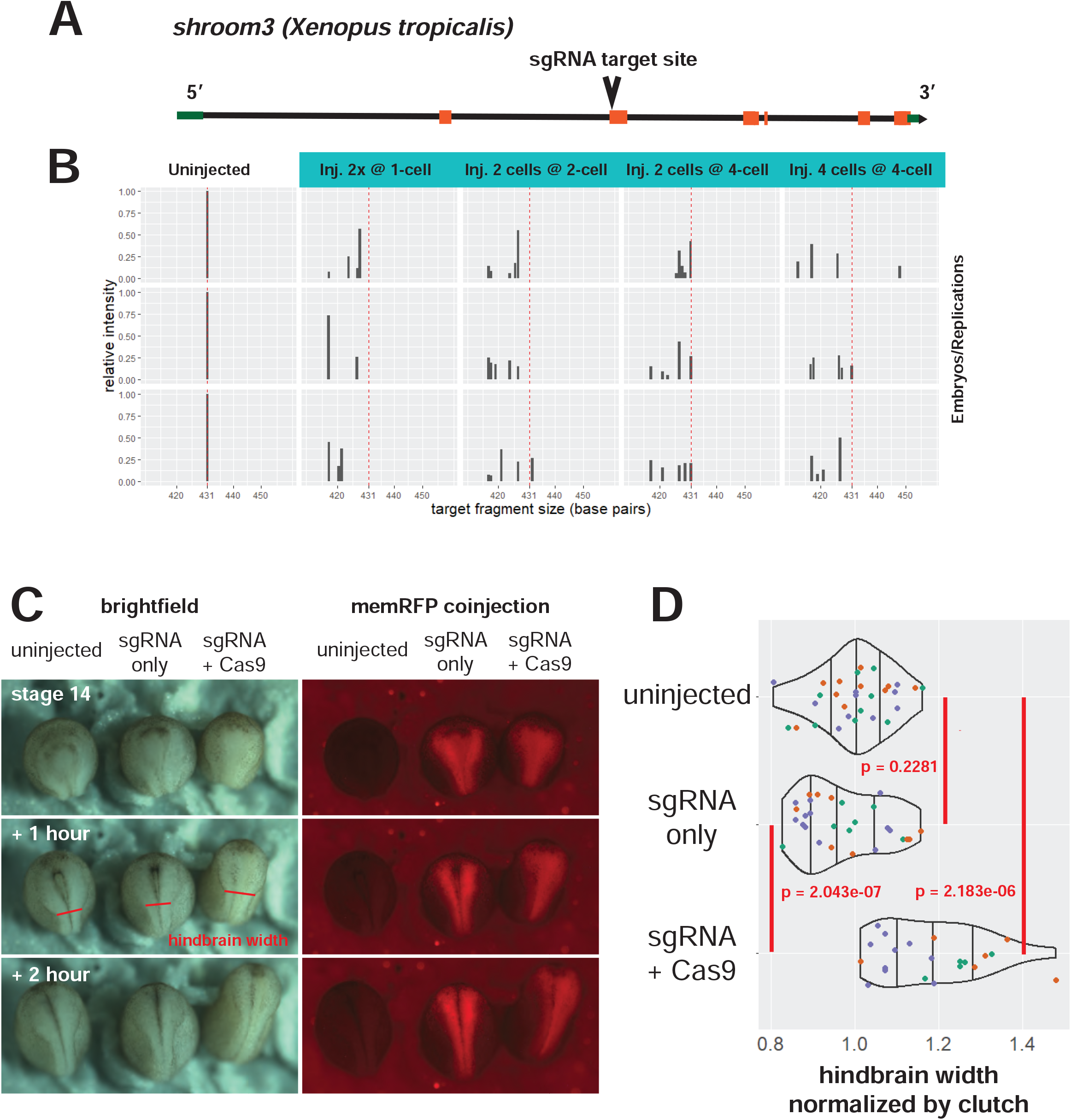
*shroom3* CRISPR validation. **A**. Gene model for *shroom3* in *Xenopus tropicalis* per Xenbase. The sgRNA was designed to target the 5’ end of the second exon. **B**. Fragment analysis to validate efficacy of sgRNA + Cas9 injections. A 431 base pair genomic fragment of *shroom3* at the sgRNA target site was generated by PCR from whole embryo lysates and subjected to capillary electrophoresis. Uninjected embryos that did not receive CRISPR reagents showed one band at 431 base pairs (left column, red dashed line). Embryos injected with *shroom3* sgRNA + Cas9 protein at various times and frequencies are displayed in right-hand columns, with the wild-type fragment size indicated by the red dashed line. Relative frequency of wild-type *shroom3* fragment size is severely reduced in injected embryos. Each plot represents one embryo. **C**. sgRNA controls. Embryos were injected with *shroom3* sgRNA alone or sgRNA plus Cas9 protein into the dorsal blastomere at the 8-cell stage, as in the imaging experiments. membrane-RFP mRNA was used as injection/lineage tracer. Red line segments indicate the posterior boundary of the hindbrain, which was used to calculate values in D. **D**. Quantification of sgRNA controls. Hindbrain widths were measured at approximately stage 17 as indicated in C. Colors of individual points indicate clutch membership of each embryo. An average width of the hindbrain of uninjected embryos of each clutch was calculated, then all embryos in each clutch had their hindbrain width calculated as a ratio of that average. Density distribution violins were then calculated based on pooled clutches. Vertical lines on density plots/violins indicate quartiles of distribution. Statistical tests were calculated using T-test.

**Methods Appendix Figure 2:**
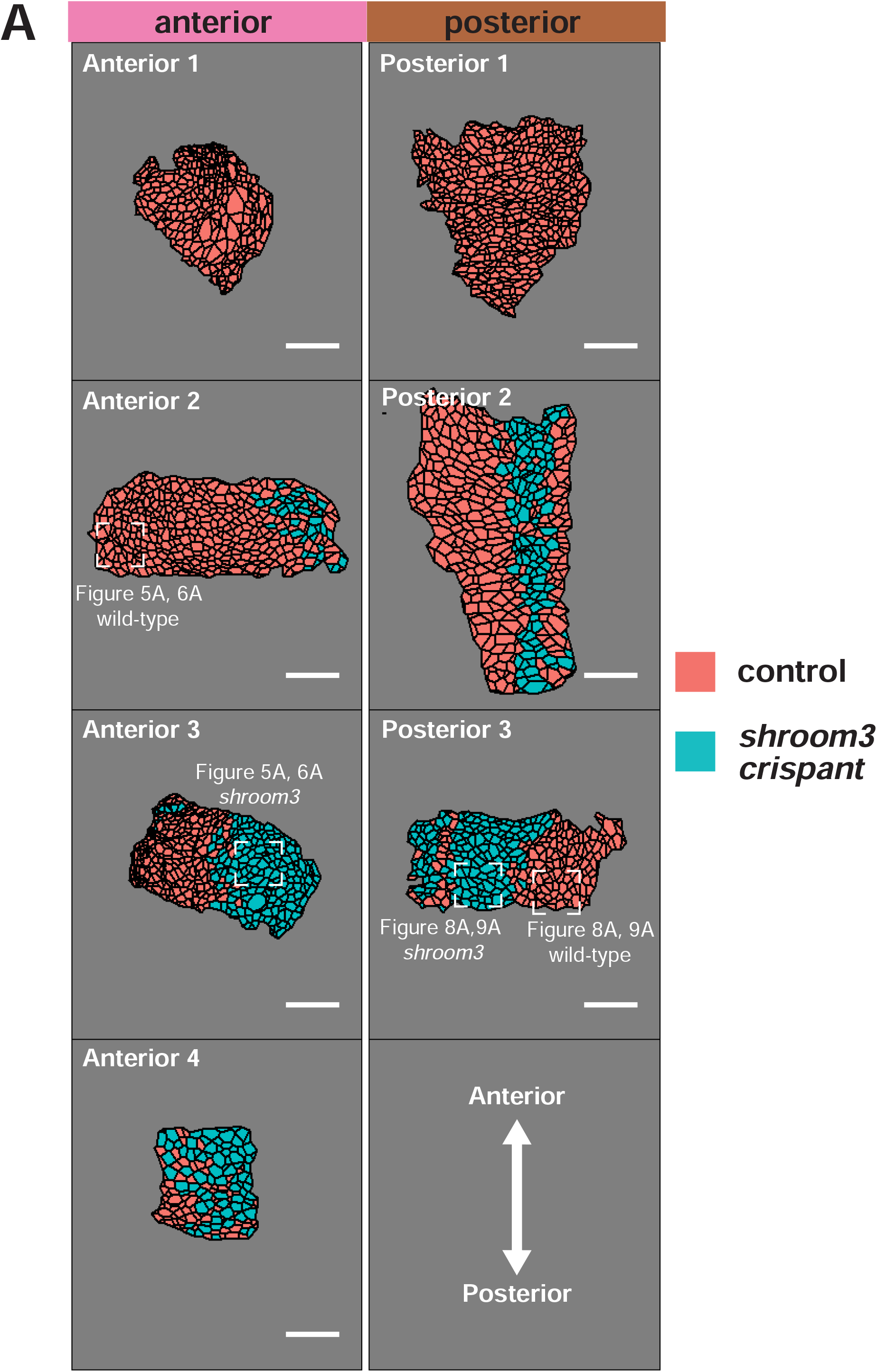
Maps of initial frames and *shroom3* crispant calls for each analyzed embryo. Representative cells in Figures 5, 6, 8, & 9 are highlighted. Scale bar = 100 microns.

**Methods Appendix Figure 3:**
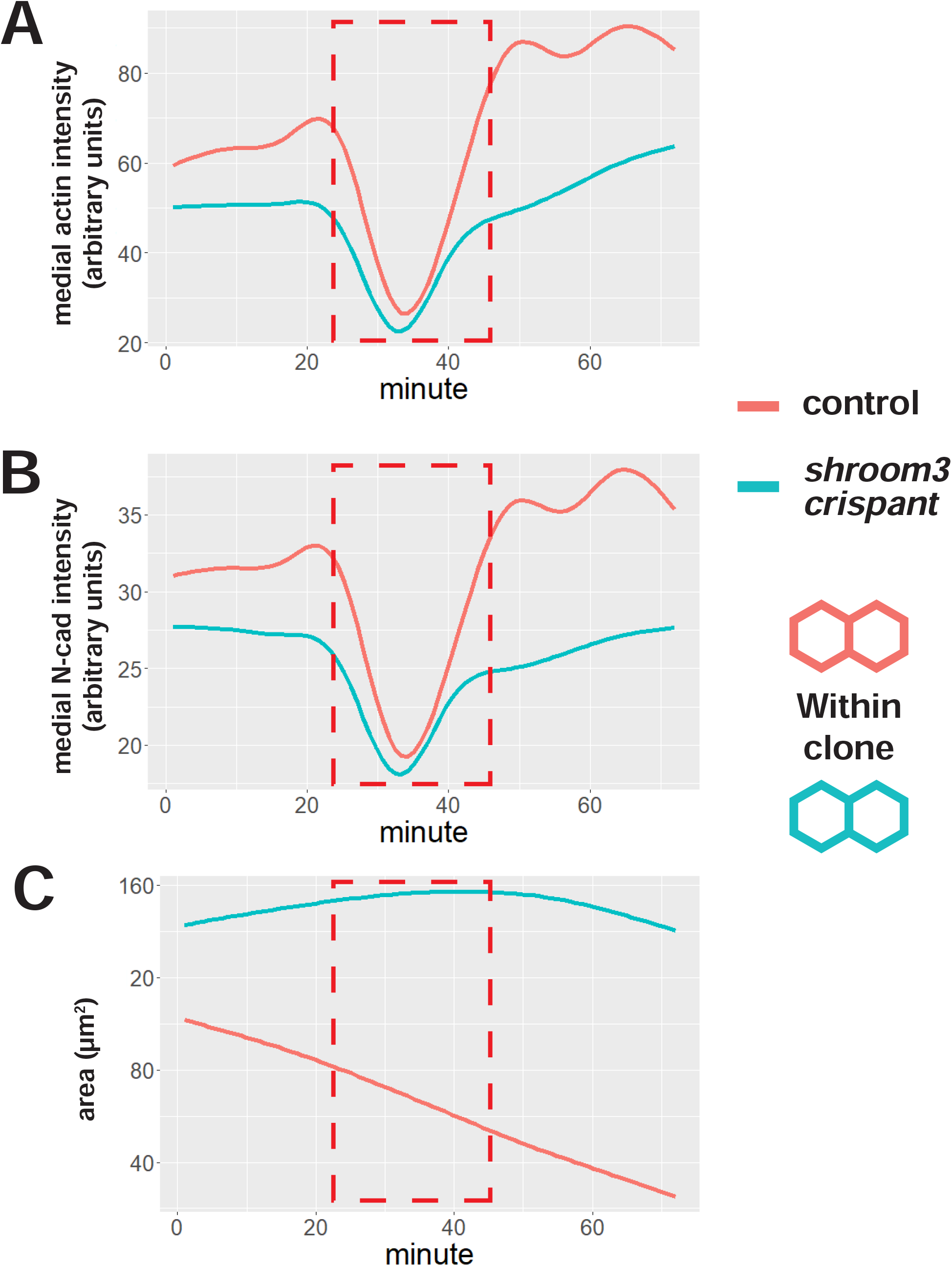
Fluorescent aberration in an anterior-imaged embryo. **A**. Smoothed mean of medial LifeAct-RFP/actin fluorescent intensity across all imaged cells across time. **B**. Smoothed mean of medial N-cadherin-GFP fluorescent intensity across all imaged cells across time. **C**. Smoothed mean of apical surface area across all imaged cells across time. Red dashed box indicates frames where fluorescence values dropped temporarily. Cells were tracked through these frames, but the frames were removed from analysis of this embryo.

**Methods Appendix Figure 4:**
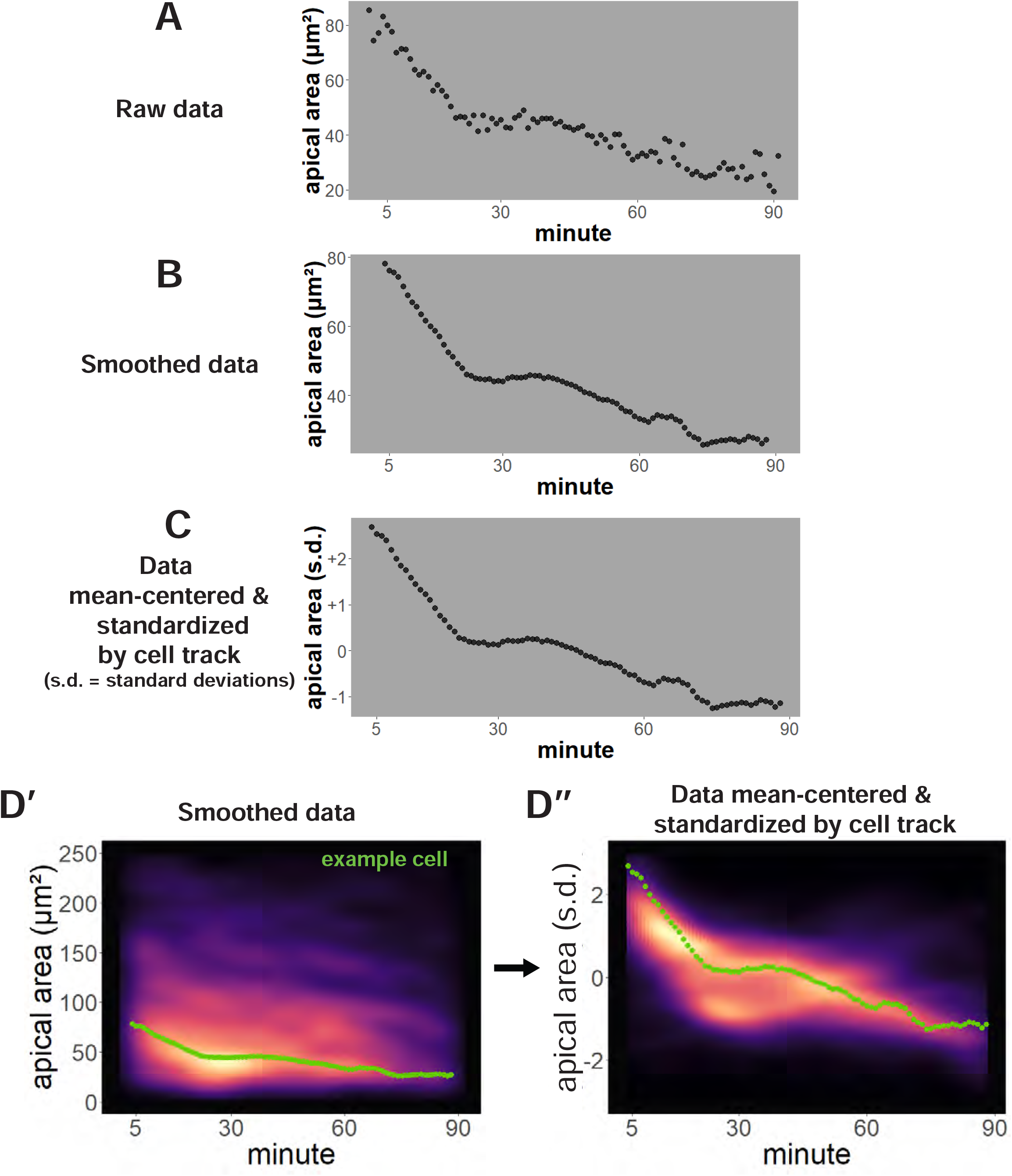
Per-cell data processing and analysis. **A**. Raw data for apical area (converted to square microns) of an individual cell over time. **B**. Apical area averaged/smoothed over 7 frames. **C**. Smoothed data after mean-centering and standardization. **D’**. Apical area (square microns) versus time of all cells from a control embryo displayed as a density plot. The green dots are from the same cell as displayed in B. **D”**. The same cells as D’ after mean-centering and standardization of each cell track. The green dots are from the same cell as displayed in C. s.d. = standard deviations.

## Notes

### Competing Interest Statement

The authors have declared no competing interest.

### Summary of Updates

This revision includes a more thorough treatment of controls used for the CRISPR experiments.

